# Normal development of the brain: a survey of joint structural-functional brain studies

**DOI:** 10.1101/2021.04.20.440563

**Authors:** Roxana Namiranian, Sahar Rahimi Malakshan, Hamid Abrishami Moghaddam, Ali Khadem, Reza Jafari

## Abstract

Joint structural-functional (S-F) developmental studies present a novel approach to address the complex neuroscience questions on how the human brain works and how it matures. Joint S-F biomarkers have the inherent potential to model effectively the brain’s maturation, fill the information gap in temporal brain atlases, and demonstrate how the brain’s performance matures during the lifespan. This review presents the current state of knowledge on heterochronous and heterogeneous development of S-F links during the maturation period. The S-F relationship has been investigated in early-matured unimodal and prolonged-matured transmodal regions of the brain using a variety of structural and functional biomarkers and data acquisition modalities. Joint S-F unimodal studies have employed auditory and visual stimuli, while the main focus of joint S-F transmodal studies has been resting-state networks and working memory. However, non-significant associations between some structural and functional biomarkers and their maturation show that designing and developing effective S-F biomarkers is still a challenge in the field. Maturational characteristics of brain asymmetries have been poorly investigated by the joint S-F studies, and the results were inconsistent with previous non-joint ones. The inherent complexity of the brain performance can be modeled using multifactorial and nonlinear techniques as promising methods to simulate the impact of age on S-F relations considering their analysis challenges.

## 1. Introduction

The human brain undergoes many structural and functional changes from infancy to adulthood. Investigating these changes with the aim of closely monitoring the normal brain development process and its structural and functional pathology is of particular importance. Study of the normal brain development process reveals the secrets of the human brain which are essential for artificial intelligence-related fields (Fan et al., 2020). A vital application of this study is predicting the risk of psychiatric or neurodevelopmental disorders. Prediction and early diagnosis of neural disorders are dependent on the detection of any deviation from the normal developmental pattern of the brain. Additionally, understanding the possible impacts of environmental factors and premature birth on brain development is important in neuroscience, and consequently many studies have been aimed to explore the influence of each factor on normal development.

Structural and functional studies use various data acquisition modalities to capture different aspects of the brain during development. Magnetic resonance imaging (MRI), as a low risk modality, plays an important role in identifying and evaluating anatomical brain changes in macroscopic and mesoscopic levels. Microstructural information of the brain is mostly acquired by diffusion-weighted imaging (DWI) (Nucifora et al., 2007)—a form of MR imaging based upon measuring the random Brownian motion of water molecules within a brain tissue — especially diffusion-tensor imaging (DTI) which detects and quantifies the anisotropy of water diffusion. The brain function development can be investigated by functional MRI (fMRI) based on the brain hemodynamic response, by electroencephalography (EEG)/ Magnetoencephalography (MEG) based on electric/magnetic field over the scalp, by functional near-infrared spectroscopy (fNIRS) based on changes in concentration of oxy/deoxygenated hemoglobin, and by positron emission tomography (PET) based on changes in metabolic processes visualized by radiotracers.

Maturation of gray matter (GM) and white matter (WM) over the course of development relies on a succession of cellular events, driven by biological clocks or entrained by the sensory inputs (Herschkowitz, 1988; I Kostović & Judaš, 2015). The age-related changes of structural features of GM (Gogtay et al., 2004; Sowell et al., 2003) and WM microstructures (Dubois et al., 2014; Lebel et al., 2019) both show heterochronous and heterogeneous inverted U-shape trajectory but with different changing rates (Dubois et al., 2014; Gilmore et al., 2018; Lebel et al., 2019). After an initial increase in GM in the first years of life, synaptic pruning (Hong et al., 2016) and scheduled modifications in size and number of glia or size of neurons (Varela-nieto et al., 2003) and myelination (Miller et al., 2012) cause nonlinear contraction of GM in late childhood and early adolescence. In contrast, mechanisms like increase in myelination and axonal diameter (Herschkowitz, 1988; I Kostović & Judaš, 2015) are the reasons for the slow linear increase in white matter volume (WMV) from birth and even beyond adolescence into middle age up to around 50 years (Hong et al., 2016; Varela-nieto et al., 2003). From an overall point of view, the sequence of the brain evolution starts with the maturation of the lower-order primary and sensory cortical regions and related connections (Dubois et al., 2016), and continues with the protracted development of higher-order cognitive regions of the brain (Baum et al., 2020; Chomiak & Hu, 2017). The hierarchal evolvement of the brain seems to be in line with energy constraints and efficient expenditure during life (Chomiak & Hu, 2017).

The graph-theory-based approaches also have demonstrated that structural covariance networks (SCNs) of gray matter volume (GMV) and cortical thickness have a hierarchically developmental approach. The primary sensorimotor SCNs are well developed by birth, while higher-order association SCNs mature and become fine-tuned later in childhood and adolescence (Zielinski et al., 2010). However, the main properties of structural and functional networks like modularity and local and global efficiency show that the organization of the neonatal brain is the same as the mature adult brain. The organization of WM networks including rich-hub property before birth is similar to that of mature networks in adulthood (Cao et al., 2016; Yap et al., 2011). The modules and hubs regions in the WM network which are in place by 2 years of age continue strengthening during development in the direction of decreasing the network segregation, and increasing its integration and efficiency (Hagmann et al., 2010).

In line with the structural networks, primary functional networks including the sensorimotor, visual, and auditory networks are pioneer networks to mature and have adult-like topologies at term age (Gilmore et al., 2018). After birth, the primary connections are refined to process the signals more efficiently in an experience-dependent process (Feldman & Brecht, 2005). Higher-order functional networks develop after birth and start with synchronization of main hub regions of default mode network (DMN), as well as language network in the first year of life (Emerson et al., 2016; Gao et al., 2015). Then other high-order networks like the salience network (SN) and the dorsal attention network (DAN) establish distributed network-like topologies by 2 years of age, and the executive control networks are the last networks to mature (Gao et al., 2015). Maturation and specialization theories, respectively suggest that emerging new behavioral abilities are associated with neuroanatomical maturation and development of interactions between brain networks (Johnson, 2001).

Joint S-F analysis during development can show how structural and functional aspects of the brain are related and how their relation matures by age. The focal point of this review is the studies aimed to answer the questions about the link between structural and functional measures of the brain during normal development based on multimodal neuroimaging data. The structural (functional) studies that use functional (structural) findings to select regions or interpret their results are based on the concept of the relation between structure and function of the brain. In addition, tracing changes in the normal relation between structure and function of the brain during development would help for early diagnosis or even prognosis of mental disorders. The measures of age-related S-F association can provide us with the brain development information, beyond what functional and structural measures alone do (Zimmermann et al., 2016). It is noteworthy that multi-modal neuroimaging studies generally outperform modality-specific studies in diagnosing the neurodevelopmental disorders (Sui et al., 2012). Several review studies focused on the relation between the structure and function of the brain (Batista-García-Ramó & Fernández-Verdecia, 2018; Rudrauf, 2014; Suárez et al., 2020) as well as the age-related changes in each structural and functional aspect (Gilmore et al., 2018; Haartsen et al., 2016; Luo et al., 2020; Oldham & Fornito, 2019; Paterson et al., 2006). However, to the best of our knowledge, no review study focused mainly on the joint S-F development of the brain. Accordingly, the studies which reported the interaction of behavioral or psychological measures with only one of the structural or functional features were excluded from this review. Normal development being the primary motivation of this review, the studies investigated the structural and functional changes during disorders, as well as those considered the development of premature subjects as a deviation from normal maturation were not our key focus.

In this review, we have categorized the joint S-F development studies in unimodal and transmodal regions. Lower-order unimodal regions of the brain, like primary and adjacent visual, auditory, somatosensory cortex receive input from a single sensory modality (Mesulam, 1998). On the other hand, higher-order transmodal regions, like DMN, integrate input information from other higher-order transmodal or downstream sectors of unimodal regions (Mesulam, 1998). As described earlier in this section, the structural and functional maturation rate of unimodal sensory regions is high in the first years of life and is prolonged in transmodal regions. **Fig. 1**, provides pie histograms versus age-range for unimodal and transmodal joint S-F development studies. The circle size illustrates the relative number of various studies in each category.

**Fig. 1.**
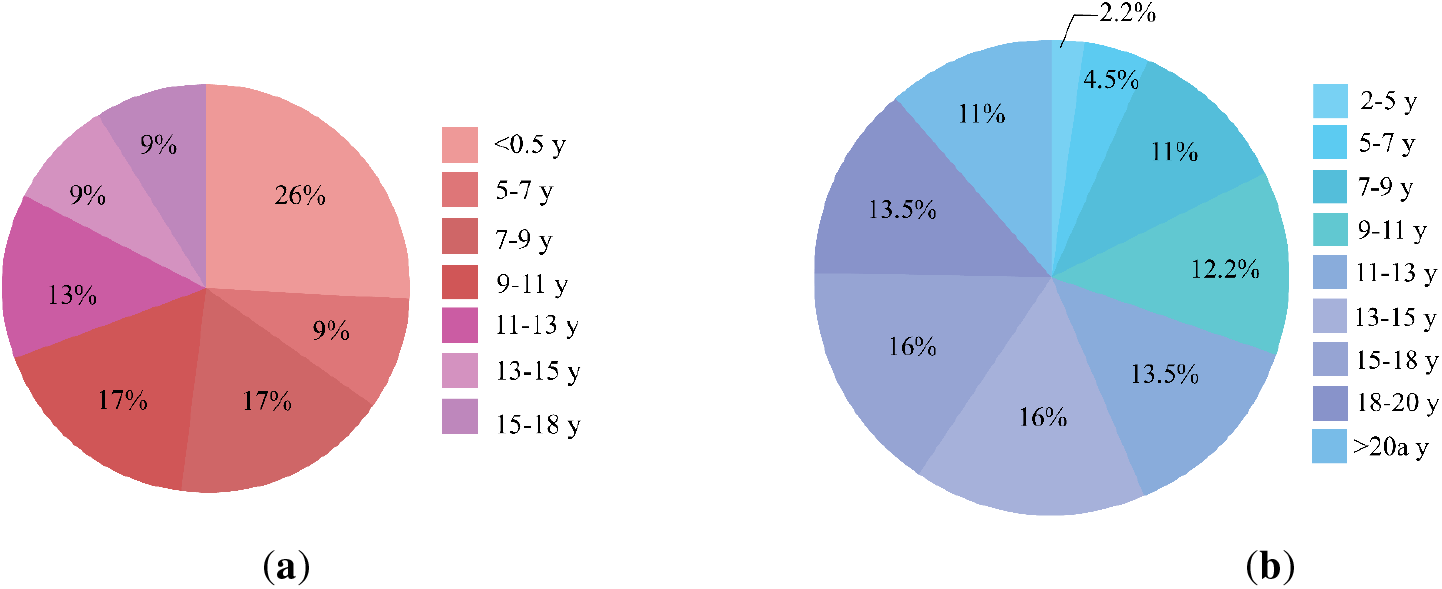
Histogram of age-ranges participated in joint S-F development studies. (**a**) lower-order unimodal regions that mature structurally and functionally earlier compared with (**b**) high-order transmodal regions with protracted development. The size of the circles corresponds to the relative number of studies in the two categories.

The remaining of the paper is organized as follows: Section 2 reviews the joint S-F studies focused on the lower-order unimodal regions of the brain. In Section 3, studies related to the higher-order transmodal regions are reviewed. Section 4, presents some main challenges in age-related joint S-F studies and draws future research directions.

## 2. S-F development in unimodal sensory regions

The human brain benefits from a well-defined modular organization in which the network modules emerge very early in life (Van Den Heuvel et al., 2015), and are remodeled and segregate increasingly during childhood and adolescence (Baum et al., 2020; Finn et al., 2015). During development, the organization of the human cerebral cortex is extending from unimodal sensory to transmodal association cortex (Huntenburg et al., 2018; Margulies et al., 2016). This section reviews the studies focusing on the age-related structural and functional changes in visual and auditory regions as summarized in **Table 1**. **Fig. 2** illustrates the unimodal regions and their developmental patterns that have been investigated in these studies.

**Fig. 2.**
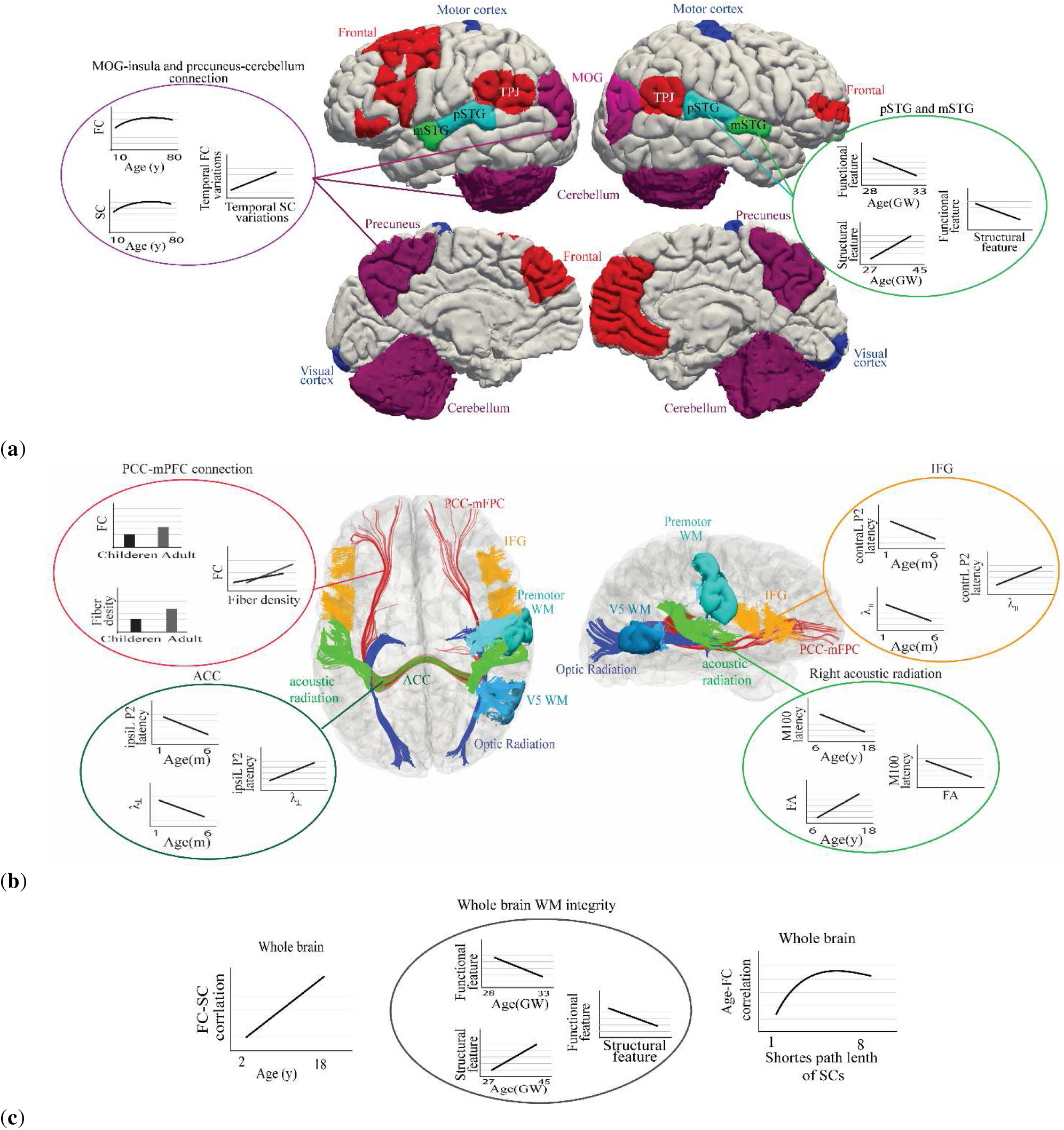
Regions with significant variation in S-F relation during development. (**a**) Cortical areas: black blue regions including motor and visual cortex with decreasing S-F relation, red regions including frontal and TPJ with increasing S-F coupling (for 8-23 years old subjects) (Baum et al., 2020). Auditory-related regions (mSTG and pSTG) in light blue and green, show related structural and functional development for infants (Daneshvarfard et al., 2020). The development pattern of SC and FC of MOG-insula and precuneus-cerebellum, in pink and purple, show increasing connections and positive S-F relation (Luo et al., 2020). (**b**) Regional WM and tracts: tracts between PCC and mPFC in red show increasing FC and fiber density during development (Supekar et al., 2010). Blue tracts show optic radiation with increasing transverse diffusivity and association with the speed of visual P1 (Dubois et al., 2008). The maturation of WM of V5 and premotor regions, in light blue, during childhood is associated with the development of visual response’s speed (Dockstader et al., 2012). Transverse diffusivity in ACC, colored in dark green, and longitudinal diffusivity in IFG, colored in orange, are correlated with the latency of left ipsiL and averaged contraL P2, respectively (Adibpour et al., 2020). S-F relation between FA of acoustic radiation, in green color, and M100 latency is significant in the right hemisphere (Roberts et al., 2009). (c) Whole brain WM and tracts: Increasing SC-FC relation during development (2-18 years) (Hagmann et al., 2010), development pattern of the WM integrity for infants (Daneshvarfard et al., 2020), and maturation of FC in indirect SCs (Roberts et al., 2009), considering the whole-brain S-F development are shown from left to right. *ACC* auditory fibers of the corpus callosum, *FC* functional connectivity, GW gestational week, *IFG* inferior frontal gyrus, *m* months, *mPFC* medial prefrontal cortex, *MOG* middle occipital gyrus, *mSTG* middle superior temporal gyrus, *PCC* posterior cingulate cortex*, pSTG* posterior superior temporal gyrus, *SC* structural connectivity, *TPJ* temporoparietal junction, *y* years, λ_⊥_ transverse diffusivity, *λ*_∥_ longitudinal diffusivity.

**Table 1.**
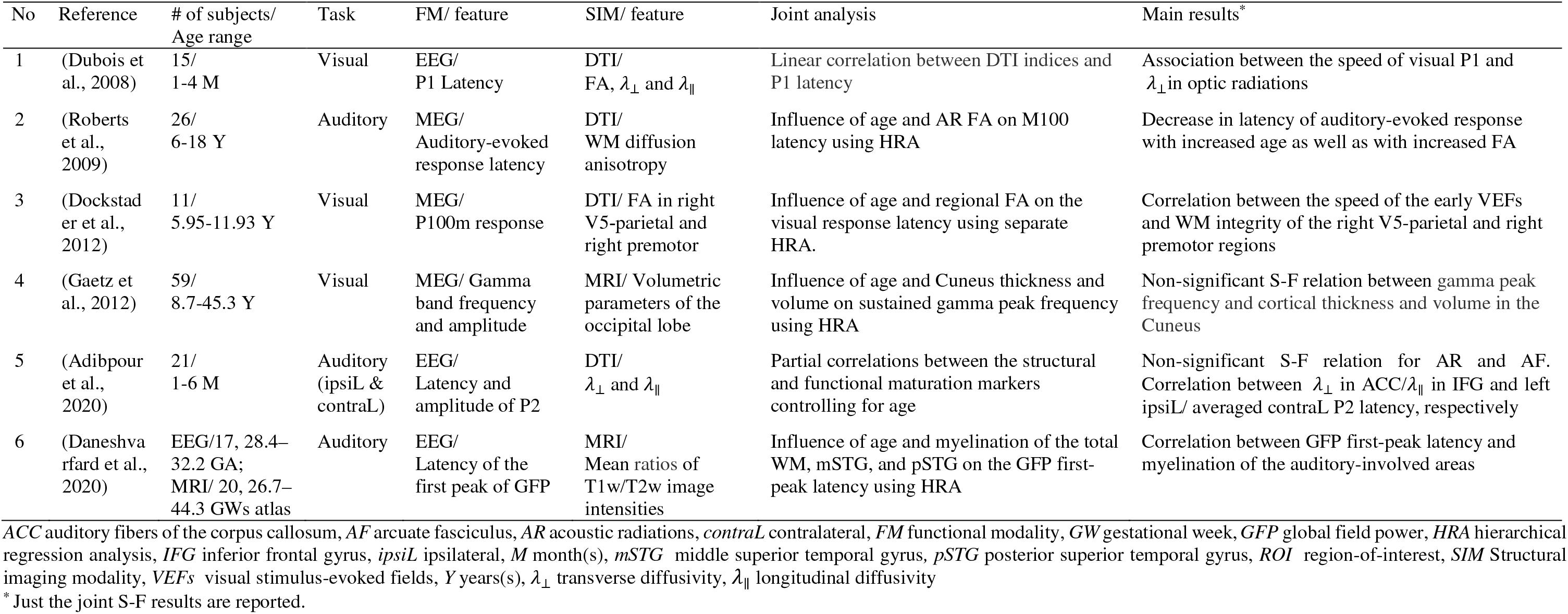
Summary of the studies focusing on joint S-F age-related changes in visual and auditory sensory regions of the human brain.

### 2.1. Visual brain areas

General development of the visual system starts very early in fetal life and continues throughout the whole lifetime. Before birth, cortico-cortical callosal and projection pathways are developed. After birth, the myelination of optic radiations starts and continues through six postnatal months, while the myelination of visual callosal fibers continues through childhood (Dubois et al., 2014). In order to investigate the maturation in projection and interhemispheric cortico-cortical pathways, Dubois et al. (Dubois et al., 2008) assessed the correlation between the conduction speed of the early visual responses, P1 peak, obtained using EEG and transverse diffusivity in optic radiation obtained using DTI during the first semester of postnatal life. Dockstader et al. (Dockstader et al., 2012) reported that the speed of early VEFs recorded by MEG in 6-to12-year-old children was inversely related to WM integrity. Their achievement was based on computing fractional anisotropy (FA) from DTI data in visual and motor association regions. It was a strong ground to show that WM fibers which connect the ocular motor areas to visual regions, transfer information that modulates the visual response. In addition to P1, P300 is appropriate for studying functional brain development over adolescence, a period when cognitive functions are still developing rapidly (Crone & Steinbeis, 2017). It is worth mentioning that both myelination of visual tracts and development of visual perception occur in early years of life. Nevertheless, the maturation of myelin’s properties continues with further developments such as increase in the fiber diameter and in compactness and cohesiveness of fiber tracts, as well as decrease in extra-axonal spaces and cerebral fluids (Beaulieu, 2002; Schmithorst et al., 2002). The authors in (Stufflebeam et al., 2008) demonstrated similar S-F relationships in adults with a negative correlation between the latency of visual P1 and WM microstructure of bilateral parietal and right lateral frontal WM. They interpreted that the speed of information transmission depends on the establishment of lipid myelin around neuronal axons and axon diameter, both of which facilitate the synchronized neuronal communication responsible for higher-order cognitive functions (Dockstader et al., 2012).

Dubois et al. (Dubois et al., 2008) measured P1 conduction velocity of visual evoked potentials (VEPs) using EEG data and transverse diffusivity in various WM tracts using DTI data from full-term infants. They demonstrated that FA and transverse diffusivity are structural markers of myelination. They related the increase in P1 conduction velocity of VEPs to maturation of optic radiations and the transfer time between hemispheres to maturation of the visual callosal fibers. Microstructural changes in these areas reflect the pattern of growth in children that continues toward adulthood (Gomez et al., 2017).

A joint S-F study (using MRI and MEG) assessed the age-related correlation between gamma band responses of the primary visual cortex and structural measures including the cortical thickness, volume and surface area of pericalcarine and cuneus (Gaetz et al., 2012). Significant correlations were observed between the age and both structural and functional measures. However, hierarchical regression analysis (HRA) demonstrated the unique effect of age on predicting the most variations of the gamma peak frequency. Also, the thickness of the cuneus described a slight percentage of the variations. Another study showed a significant correlation between gamma-band oscillations in human participants and the surface area of the visual cortex without considering the age. The authors found no significant change between age and functional measure may be due to the relatively narrow age range (Schwarzkopf et al., 2012).

### 2.2. Auditory brain areas

Auditory information from the environment is conveyed to the brain through the primary auditory pathways (messages from cochlea) and non-primary ones called reticular sensory pathways (from all types of sensory messages) (Pujol et al., 1998). The development of human cochlea is completed by birth, while the brain’s auditory pathways and centers show progressive development from the brain stem to the auditory cortex in response to the stimulation (Litovsky, 2015; Pujol et al., 1998). It has been shown that relocation of thalamic afferents from the transient subplate zone to the cortical plate results in the emergence of cortical evoked responses to sounds (Graziani et al., 1974). The ability to perceive a change in the frequency of tonal stimuli undergoes maturation starting from three months of postnatal age, and cortical auditory evoked potentials (CAEPs) could be used to understand the neural substrates of auditory perception (Cone & Whitaker, 2013). There are invaluable studies regarding CAEPs from early infancy to adulthood, which have generally stated a decrease in latencies of CAEP components with age (Lippé et al., 2009; Wunderlich & Cone-Wesson, 2006).

Roberts et al., (Roberts et al., 2009) discovered a decrease in latency of auditory responses in the right superior temporal gyrus (STG) associated with the maturation of acoustic radiations of the auditory pathway, which was measured by FA in adolescents and children. However, for the left hemisphere, marginally significant developmental changes of functional measure and no significant S-F relation were supposed to be due to an insufficient number of subjects. In contrast, by studying full-term infants, Adibpour et al. (Adibpour et al., 2020), observed a correlation between the inter-individual variability in P2 responses and microstructural properties of GM maturation in the inferior frontal region but not in acoustic radiations. However, the functional lateralization of P2 responses was not related to microstructural asymmetries of optic radiation between two hemispheres. In another study, Daneshvarfard et al. (Daneshvarfard et al., 2020) related developmental changes of CAEPs, characterized as the decrease of the first-peak latency of the global field power (GFP), to the structural myelination index (mean ratios of T1w/T2w image intensities) of auditory-involved areas extracted from MR images of the same age infants. Nevertheless, neither the structural maturation index nor age contributed to significant additional variance in the GFP first-peak latency after accounting for the variance associated with the other parameter.

## 3. S-F development in transmodal regions

By integrating information from different higher-order regions and functionally specialized modules, transmodal regions organize the human brain as large-scale neurocognitive networks and enable humans to outperform other species in high-order tasks like language, memory-executive functions, and cognition. (Mesulam, 1998). The prolonged development of higher-order cognitive and executive functions and functional specialization in the related areas are preceded by the targeted activity-dependent myelination and synaptic remodeling in the transmodal association cortex. Structural and functional maturation of higher-order association cortex and their connections expand until early adulthood. In this section, we will review the studies on development of transmodal areas in resting state (RS) and during working memory. **Table 2** resumes the analyses and results of the joint S-F studies, reviewed in this section. **Fig. 2** illustrates the transmodal regions and their developmental patterns that have been documented in the studies indicated in **Table 2.**

**Table 2.**
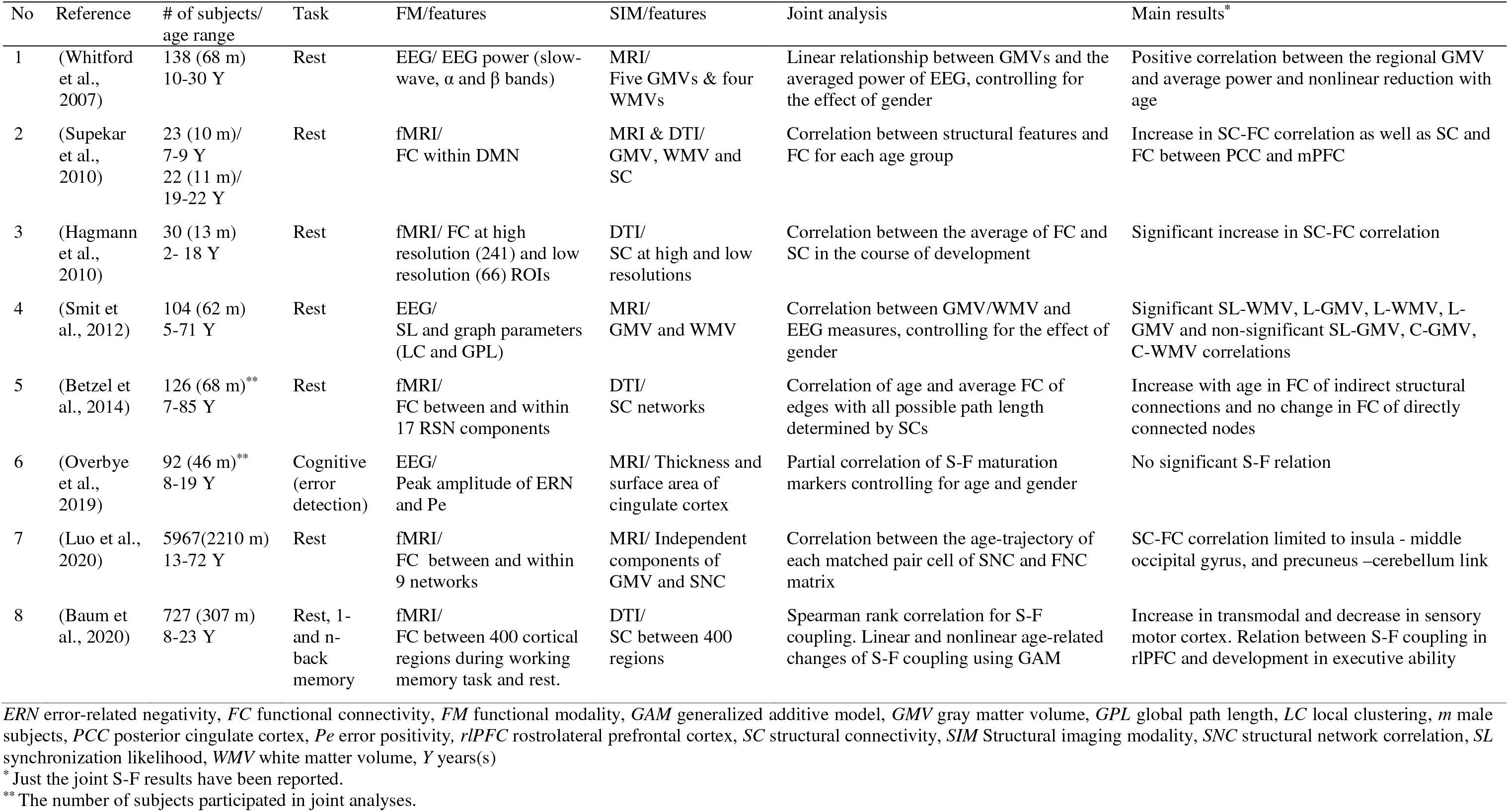
Summary of the studies focusing on joint S-F age-related changes in transmodal regions of the human brain.

### 3.1. RS studies

The RS studies have investigated the simultaneous activation of various regions of the brain as resting-state networks (RSNs). The brain networks are active during rest and form the intrinsic functional organization of the brain. The emergence of RSNs starts with primary sensory-motor and auditory networks at term age, followed by the visual network and then networks related to higher-order processing like attention and DMN and ends by the executive control-related networks. The changes of higher-order networks extend during development and lifespan (Gilmore et al., 2018). In addition to the different developmental courses of RSNs and their functional variety, the feasibility of their recording in prenatal age and during lifespan makes them attractive in age-related functional and S-F studies.

DMN is a core network exhibiting high metabolism, in social cognitive and self-monitoring functions (Spreng et al., 2009), and during RS and low activity as the brain engages in a task (Immordino-Yang et al., 2012). It has the potential to render biomarkers for estimating age, as well as psychiatric and neurological disorders and environmental advertises (Rebello et al., 2018). The association of DMN in various states and tasks makes it the center of attention in many cognitive and most RS studies. By a way of illustration, precuneus as the hub region of DMN shows activity during RS and other attention-demanding tasks like memory retrieval, reward monitoring, and emotion processing (Cavanna & Trimble, 2006).

S-F age-related studies, especially region-wise ones, demonstrate that by the early stage of life, the composition of most nodes and hubs of the brain matures like adults (Hagmann et al., 2010; Supekar et al., 2010), while the structural connectivity (SC) and FC between these regions mature heterogeneously. The posterior cingulate cortex (PCC) and medial prefrontal cortex (mPFC) have more significant GMVs in childhood, which suggests great synaptic connectivity. The developmental trajectory of these nodes continues with synaptic pruning and GMV reduction (Supekar et al., 2010; Thompson et al., 2005). These two nodes also have weaker FC in childhood versus adulthood, evaluated by partial correlation after controlling for the influence of other nodes of the DMN and non-DMN brain networks in childhood. The SC-FC significant positive trend between PCC and mPFC emerging during development suggests the growing stability during adolescence (Supekar et al., 2010).

On the contrary, the GMV of some nodes such as the medial temporal lobe (MTL) do not change significantly with age, but their connections with other nodes like PCC change differently in each hemisphere during development. The PCC-right MTL structural connection changes non-significantly while an increase in the fiber density as well as FA in the left cingulum (hippocampus), strengthens PCC-left MTL structural connection in the course of development. As the structural and functional maturation patterns of different brain connections indicated, the early maturation of FC is followed by the emergence of PCC-left MTL structural connections (Supekar et al., 2010), insula and middle occipital gyrus as well as precuneus and cerebellum (Luo et al., 2020). In adulthood, by taking age as a covariate, the FA and FC of cingulum are correlated positively, while none of these measures was associated with age (Van den Heuvel et al., 2008). Besides, long-range connections against short pathways afford delayed structural and functional maturation (Hagmann et al., 2010; Supekar et al., 2010). Notably, the high FC in any age-stage with sparse direct SCs or before the maturation of the related SCs is considered to be mediated through the indirect structural link (Fjell et al., 2017; Honey et al., 2009; Luo et al., 2020). In childhood, for example, despite of weak PCC-left MTL SC, there is adult-like level of PCC-left MTL FC as well as robust interhemispheric FC between right and left MTL (Stark et al., 2009; Supekar et al., 2010).

The whole-brain and high-resolution S-F studies demonstrated how age-related changes of short and long connections affect the brain function (Betzel et al., 2014; Hagmann et al., 2010). By the age of 2 years, direct and short efficient pathways whose structural and functional relations change weakly with age have formed the connections between nodes. Afterward, long pathways (Hagmann et al., 2010) and the indirect structural connections (Betzel et al., 2014) mature and the neural synchrony as a consequence of this maturation causes stronger FC and SC and SC-FC coupling. As a result of protracted SC and FC, increasing efficiency and decreasing clustering (Hagmann et al., 2010) or RSNs modularity (Betzel et al., 2014) as well as declining the randomness of the connectivity graph (Smit et al., 2012) betide by the course of time during development.

The age-trajectory of FC has a region-wise pattern during teenage and adult lifespan FCs between RSNs change linearly while FCs between frontal and temporal areas are the examples of quadratic shaped age-trajectory (Betzel et al., 2014; Fjell et al., 2017; Luo et al., 2020). The difference between the linear pattern of the FC changes and evolution of SC, which changes in an inverted U-shaped pattern (Fjell et al., 2017; Luo et al., 2020), stems from their different origins. The density of GMV (Terribilli et al., 2011) affects FC changes, while demyelination, gliosis, Wallerian degeneration and reduction of the number of fiber tracts can be mentioned as the sources of SC changes in WM (Burzynska et al., 2010). In S-F coupling, the GM measurements were not as predictable as the properties of the functional network based on glucose consumption measured by PET (Romero-Garcia et al., 2014). However, investigating the relationship between functional and structural cortical connectivity networks with the same age-course patterns revealed the correlation between structural and functional components in higher-order areas related to face and emotion recognition during adulthood (Luo et al., 2020).

EEG signals in S-F studies can provide information about the S-F relation within each frequency band of neural oscillations. The maturation of RS-FC assessed by EEG coincides with developing thalamo-cortical and cortico-cortical connections (Jones, 2009; Ivica Kostović & Judaš, 2010; Lamblin et al., 1999). A joint S-F investigation demonstrated brain structure and microstructures are negatively correlated with FC in high-frequency (beta, gamma) bands, while FC in lower frequency (delta, theta, alpha) bands is positively correlated with the brain maturation (Birca et al., 2016). Another study (Whitford et al., 2007) demonstrated that the reduction of synapses in GM via protracted maturation of frontal and parietal lobes gives rise to a reduction in absolute EEG power in the slow-wave band. This relation in slow-wave is stronger and spatially more homogenous than in beta and alpha bands. This frequency-wise structural and functional association trend reveals the origin of different frequency oscillations and aspects of information flow. Low-frequency components of EEG signals are correlated with highly synchronous local neuronal activity between cortical pyramidal neurons. Thus, during puberty, major elimination of neurons or neuropils co-occurs with a decrease in the amplitude and power of low-frequency EEG signal components. In addition, the increase in functional integration and information flow between distant cortical regions during development (Fair et al., 2008, 2009; Smyser et al., 2010) is thought to be mediated via low-frequency oscillations (Canolty & Knight, 2010).

On the other hand, high-frequency components of EEG signals are thought to support the local processing of stimuli segregation (Canolty & Knight, 2010) and asynchronous activity between cortical pyramidal neurons and the inability to reset their phases (Varela et al., 2001). The S-F studies demonstrate that increased high-frequency FC is associated with the immature structure of the brain in the early stages of life (Birca et al., 2016). However high-frequency signals are not correlated with the reduction of GMV and cortical synapses during adolescence as opposed to low-frequency signals (Whitford et al., 2007).

In joint S-F investigations, selecting structural and functional markers to be age predictive and have a correlation with each other is a challenging topic. The relationship between EEG power and GMV (Whitford et al., 2007), as well as positive correlation between brain activity (measured by EEG amplitude and spontaneous brain activity) and cortical surface areas and volumes, signify that the reduction in GMV gives rise to a reduction in the amplitude of EEG activity and absolute EEG power (Benders et al., 2015). In addition, the brain activity at the birth time is a marker of subsequent faster growth of total cerebral brain volumes, especially for basal ganglia/thalami at term equivalent age (Benders et al., 2015). On the other hand, as mentioned previously, some studies suggested that GM changes alone cannot explain brain’s functional changes (Onoda et al., 2012; Romero-Garcia et al., 2014). The power spectral features like power spectral density (PSD) and amplitude envelope correlation (AEC) are better predictive features for adult age compared to functional features that measure the similarity of the connectivity profiles across frequency bands like inter-layer coupling. However, power spectral features of the MEG signal are not as predictive as the structural features (T1-weighted voxel intensity of cortical, subcortical, WM and CSF segments) (Xifra-Porxas et al., 2019).

### 3.2. Memory

The growth of cognition from infancy to adulthood is associated with the development of information processing factors including the working memory, control of attention and inhibition of proponent schemes and self-regulation. The working memory maturation is a key aspect of human cognitive development (Camos & Barrouillet, 2018; Cowan, 2016). Working memory is a limited amount of information that is temporarily accessible for cognitive tasks such as language comprehension and production, reasoning, problem-solving, and decision-making (Cowan, 2016; Simms et al., 2018).

The relation between structure and function of the brain in transmodal regions of frontoparietal network (FPN) and DMN (posterior cingulate) during a working memory task predicts executive performance (Baum et al., 2020). In line with the strengthening of S-F coupling, the increasing segregation of these networks and intermodular integration during adolescence indicate an increase in executive efficiency due to network specialization and decline in competitive intervention between them (Baum et al., 2017; Fornito et al., 2012; Van Den Heuvel et al., 2009). Moreover, the right rostrolateral prefrontal cortex, which is responsible for information management between FPN and dorsal attention network (Dixon et al., 2018), promotes age-related changes in executive function from childhood through adulthood (Baum et al., 2020). Considering the rise in coupling through transmodal regions, age-related reductions in coupling within unimodal sensory regions are noticeable during 1-back and 2-back memory tasks (Baum et al., 2020).

P300 is an essential component of EEG signals that is evoked when recognition of stimulus activates memory operations. The P300 subcomponent P3b related to subsequent memory processing appears in response to infrequent target stimuli, among repeated and frequent stimuli, while P3a appears when a distractor or novel non-repeated stimulus emerges among target and frequent stimuli and during task processing. P3a reaches maturity in late childhood, while P3b matures later in adolescence (Kaufmann et al., 2018; Overbye et al., 2018). Overbye et al. (Overbye et al., 2018) demonstrated that the cortical surface area in the left temporal lobe increases with the strength of P3b in 8-19 years old subjects and indicates the developing attentional performance.

Another cognitive function that evolves as the brain matures is error processing. Error positivity (Pe), a P300 response to internal error detection, and error-related negativity (ERN), a negative deflection in EEG signal following an incorrect response, are indicators of brain maturation in error processing. Although these signals emerge from the cingulate cortex, the main challenges to find S-F relation were determining appropriate age span and anatomical markers of the cingulate cortex. The peak amplitude of ERN and cortical thickness of the cingulate cortex as the markers of functional and structural maturation had no significant relationship (Overbye et al., 2019).

## 4. Discussion and future lines of research

Joint studies investigating correlated structural and functional features to model the brain’s development process were the main focus of this review article. As previous sections revealed, the longitudinal relation between brain structure and function is not straightforward and readily predictable based on simple hypotheses, conventional features, and experimental investigations. One of the reasons is the brain’s intrinsic plasticity, which allows, apart from age, other internal and external factors like genetic, gender, education, and gestational experience to influence the structures and functions of the brain and their association during development. The neuroplasticity of the brain has a heterogeneous critical period timing, emerged by the maturation of the gamma-Aminobutyric acid (GABA) circuit and limited by molecular brakes which yield to the stability of the brain after normal development (Kolb et al., 2017; Takesian & Hensch, 2013). From the perspective of cerebral plasticity, in this review, we have focused on the lifelong functional and anatomical modifications in the normal developmental trajectory, also known as developmental plasticity (Ismail et al., 2017).

A major outcome of joint studies would be creating S-F atlases of normal brain development. Principally, these atlases can perform better than simple structural or functional ones in predicting cognitive and behavioral diseases. To the best of our knowledge, few works have been concerned by the clinical applications of the created joint atlases and their advantages with respect to the previous non-joint ones. Such a clinical study requires longitudinal data by following up the subjects under study, which is a challenging issue. The future research lines and steps to achieve better applicable joint developmental atlases for the prognosis of the brain disorders are addressed in this section and summarized in **Fig. 3**. Before being able to support the psychiatrists in diagnosis and medication, joint developmental studies must fulfill a number of prerequisites that are discussed hereafter. In addition to the joint S-F biomarkers, which model normal development, the joint S-F biomarkers responsive to disorders can provide useful knowledge to create atlases aimed at early detection of disorders (**Fig. 3(a)**).

**Fig. 3.**
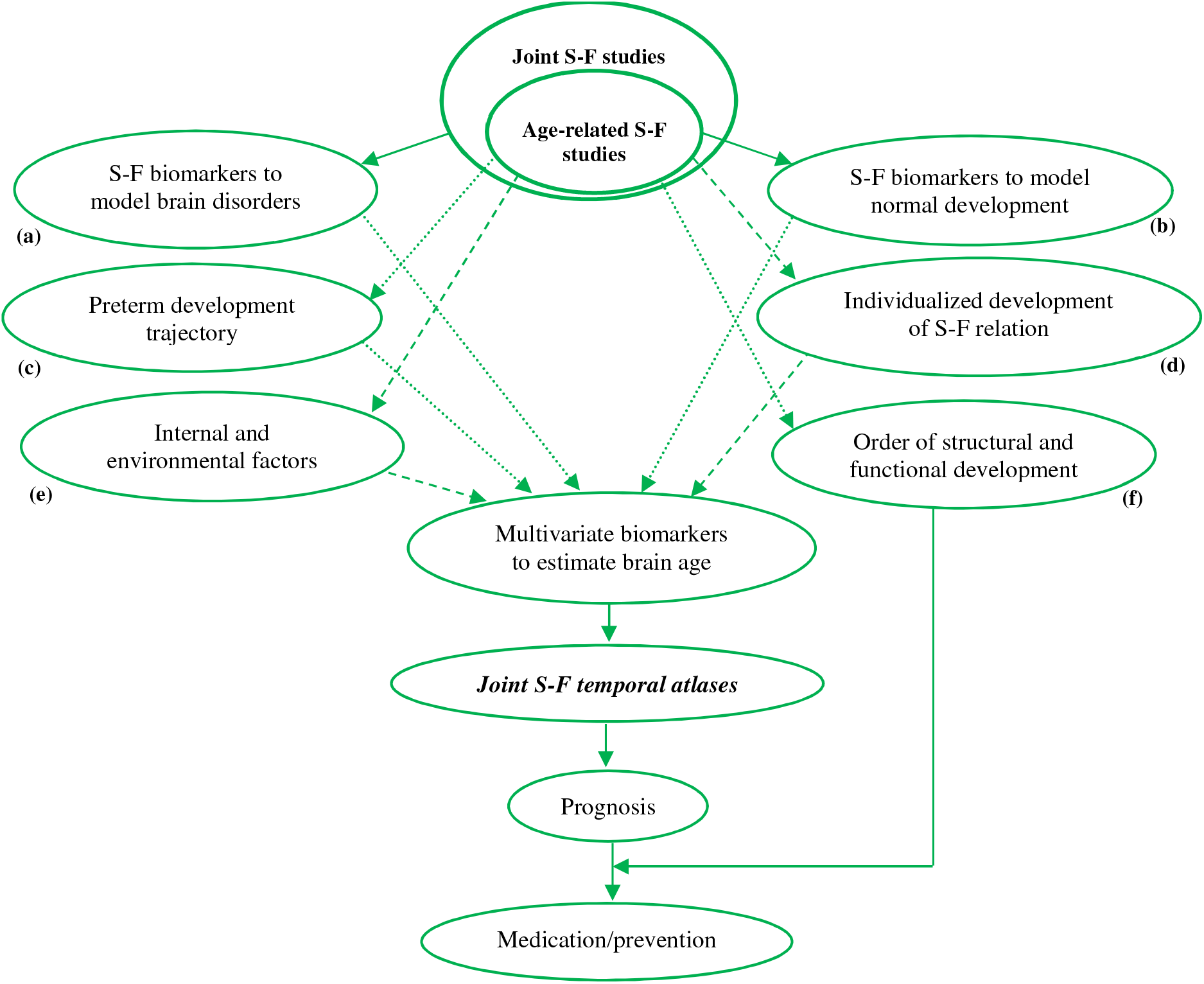
Overall view of current and future research lines and steps. This diagram shows how age-related S-F studies can lead to medication and prevention of brain disorders. Solid arrows represent the non-controversial results of studies. Dotted arrows represent the lines of research in age-related S-F studies with little attention or discordant results, suggesting that further studies are needed. Dashed arrows refer to further areas of future research in S-F developmental studies. Joint S-F studies help prognosis by introducing the multivariate biomarkers and mapping their normal maturation. What are the steps forehead to achieve suitable biomarkers and joint S-F temporal atlases? Joint S-F biomarkers, which provide complementary information to brain disorders vs. structural and functional biomarkers (**a**), as well as those joint S-F biomarkers that provide complementary information to model brain maturation (**b**) are necessary to select suitable multivariate biomarkers. An accurate developmental atlas is needed to determine if the preterm development is a possible normal maturation trajectory or not? (**c**) Inter-subject differences of S-F maturation trajectories (**d**), and internal and external effective factors on joint S-F development (**e**) are important issues to map the normal maturation of joint S-F relation of the brain. Structural and functional relation and how their maturations affect each other can help in prediction and medication of brain disorders (**f**).

The purpose of coupled S-F development studies is to show the age-related changes that occur in the brain, beyond each modality alone. Some age-related changes can be revealed using only joint S-F analysis (Supekar et al., 2010; Van den Heuvel et al., 2008; Zimmermann et al., 2016) (**Fig. 3(b)**), like growing SC-FC coupling in bi-lateral precentral, superior temporal and left entorhinal or decreasing SC-FC coupling with age in left inferior temporal, bilateral parsorbitalis, and right parstriangularis. Nevertheless, the SC-FC correlation cannot reflect all the structural and functional variations with age in DMN hub regions like precuneus, inferior parietal and left posterior cingulate where SC or FC have stronger age-related changes than coupled measures. Investigating the link between the S-F coupling and the evolution of structural and functional changes demonstrated that structural and functional results cannot be excluded from joint S-F findings, and vice versa (Baum et al., 2020). The increasing number of features in joint S-F developmental studies arises a challenging problem known as curse of dimensionality. Despite providing complementary information, higher dimension of joint S-F features makes their clinical usage (e.g. in prediction of diseases) questionable. Using dimensionality reduction strategies and accessing larger datasets are the eventual solutions to this problem. Fortunately, different projects, such as the human connectome project (HCP), have been launched in order to collect more accessible data for neuroscience research. To estimate the normal developmental model and introduce new informative biomarkers, increasing the dataset size is favored over dimension reduction techniques.

One method for developing a joint S-F temporal atlas is to use age-related changes for each structural, functional, and S-F feature and combine them. The other applicable and ideal solution, however, is to use suitable informative biomarkers to construct a joint S-F atlas. Some of the previous studies failed to show a significant correlation between structural and functional development using well-known features. Such features seem not to be specific to model the dependencies between the structure and function of the brain during development. Recently, few researches have been conducted on correlated combinatorial features as indicators for brain age (Xifra-Porxas et al., 2019) or pathologies and diagnosing disorders (Li et al., 2020; Sui et al., 2012), using advanced feature extraction methods like convolutional neural network or other nonlinear methods. This new approach promises new achievements in this field and may introduce new multivariate-multimodal features, which better describe the evolution of brain structure or function during development. The clinical and investigational use of such multivariate-multimodal biomarkers is another significant challenge. These not very well-known biomarkers must be analyzed and interpreted based on their incorporating conventional structural and functional features.

It is noteworthy that investigating the normal brain’s maturation before birth based on preterm is a challenging issue (**Fig. 3(c)**). On the one hand, some studies considered preterm born neonates as a slight deviation from normal development (Adibpour et al., 2020; Benders et al., 2015; Daneshvarfard et al., 2020) and reported their age-related changes as the evolution of the brain before birth. On the other hand, few researches tried to reveal their different structural and functional patterns in comparison with terms (Baldoli et al., 2015; Barnes-Davis et al., 2020; Rogers et al., 2018). More works will need to be done to determine the evolution of brain S-F coupling after birth, and how preterm birth may deviate natural brain maturation. Are the reported deviations just temporary and negligible changes that will be compensated by the brain’s plasticity nature? Or will preterm birth affect the brain and behavior in the future? Do these explanations help to decide whether preterm birth is an important factor in prognosis or not? The multimodal longitudinal dataset of preterm born subjects in different age stages is ideal for such study.

Another important issue for selecting biomarkers and joint S-F temporal atlas is the S-F coupling individualization (**Fig. 3(d)**). The FC fingerprinting has been observed in RS-fMRI studies, implying that the connectome backbone and its stability act as brain-based fingerprints that individualize in different conditions. (Finn et al., 2015; Onoda et al., 2012). Investigating the association of the functional individualization and its stability with both genetic and the structures of the brain during development are completely new topics valuable for understanding new aspects of the brain and its age-related changes. This combination of findings is a critical support to select appropriate biomarkers.

Despite age-related changes of WM and GM, alteration of other factors may affect the functional changes during development and aging. For example, during the developmental period, GABAergic neurotransmission develops as well as structural measures and co-occurs with the modifications in excitatory neurotransmission (Kilb, 2012). The alteration of the GABAergic system causes increased synchrony in the gamma band frequency (Hashimoto et al., 2009), which is correlated positively with hemodynamic response of the brain in fMRI signals (Niessing et al., 2005). As well, Xifra et al. mentioned the possible influence of local GABA inhibitory function on age-related changes of beta power in the adult MEG signal (Xifra-Porxas et al., 2019). Therefore, in joint S-F studies, as suggested in (Hagmann et al., 2010), to model the structural and functional relation, it is necessary to consider chemical phenomena or other factors that contribute to the acquired functional signals. To establish a specific biomarker for the joint S-F temporal atlas, it is important to evaluate and control for other factors during growth that have an undeniable impact on the evolution of the brain. An equally important purpose of such comprehensive studies is controlling the effective factors on the plasticity of the brain for medication or compensating the destructive effects as early as possible (Ismail et al., 2017; Kolb et al., 2017) (**Fig. 3(e)**).

Furthermore, how the structural and functional properties of the brain affect each other and which ones are predictive of the others is the key issue of the joint S-F studies and for treatment and prevention of diseases (**Fig. 3(f)**). The dominant hypothesis in most S-F investigations is that the function of the brain can be determined by the structural properties of the brain (Bullmore & Sporns, 2009). Likewise, the developmental studies are focused on how the neural function is related to the underlying structural connections (Hagmann et al., 2010; Honey et al., 2009; Van den Heuvel et al., 2008; Zimmermann et al., 2016). However, use-dependent synaptic pruning, plasticity and adaptive changes of the brain’s system during development confirm the structures of the brain can also change as the result of neural activity (Feldman & Brecht, 2005; Mount & Monje, 2017). Some computational studies converge to adaptive brain network model based on feedback loop to model the cooperative relation between the structure and function during maturation (Millán et al., 2019). The joint S-F developmental studies considered timing of maturation pattern of anatomy and function and reported inconsistent results. Leading maturation pattern of functional connections during life span can show its effect on lagged maturation of structure (Luo et al., 2020). Conversely, reduction of SC with age leads to the elimination of path efficiency, which results in weak signal transmission and consequently, age-related FC reduction (Betzel et al., 2014). While, concurrent maturation for each lobe is also reported (Whitford et al., 2007). However, according to (Baum et al., 2020) longitudinal imaging for several close age stages is essential to conclude about the lead-lag relation.

Investigating the laterality of brain development is another subject of joint S-F development studies. Brain lateralization starts from the fetus and continues during development and is known as a maturation index for networks like language-related networks (Bisiacchi & Cainelli, 2021; Dubois et al., 2014; Güntürkün et al., 2020). While the structural asymmetry and functional lateralization are supposed to be related, the development of joint S-F laterality demands more study. The open question in joint S-F development is how S-F lateralization changes during the life span. Several joint studies investigated the lateralized pattern of brain structure and/or function (Dubois et al., 2014; Gaetz et al., 2012; Hagmann et al., 2010; Supekar et al., 2010), while studies on development of S-F lateralization are so limited. As previously stated, Roberts et al. (Roberts et al., 2009) demonstrated S-F relation between the M100 latency and maturation of acoustic radiation is significant only in right hemisphere. They attributed the non-significant S-F relation in the left hemisphere to the inferior number of subjects and suggested further investigation. Other studies like (Adibpour et al., 2020; Supekar et al., 2010) did not find any lateralized pattern for S-F relations. Adibpour et al. (Adibpour et al., 2020) claim that selecting related structural and functional features is crucial for determining joint S-F relationships, especially in infants. The PCC-left MTL SC changes significantly during development, while there is adult-like FC in childhood. The lateralized development of SC and FC of PCC-MTL connection can be interpreted as an increasing S-F association in the left hemisphere, while S-F analysis shows no significant S-F relationship for this connection (Supekar et al., 2010). In order to use laterality as a promising biomarker for development in joint S-F temporal atlases, further research is needed to find lateralized and related structural and functional biomarkers (Adibpour et al., 2020; Supekar et al., 2010) and the age-related changes of their joint relation (Baum et al., 2020).

There are some methodological pinpoints worth mentioning. A small number of studies have investigated a nonlinear association between S-F biomarkers and age (Baum et al., 2020), as outlined in **Table 1** and **2**. The future trend will be to pursue more accurate, complex relationships that are capable of modeling comprehensive age-related brain changes. Computing FC as a partial correlation controlling for the influence of other confounding nodes and networks is needed to be considered to obtain more reliable results in FC (Supekar et al., 2010; Van den Heuvel et al., 2008). Dynamic FC, as a promising biomarker for neurodegenerative conditions (Filippi et al., 2019) and complex brain changes during the lifespan (Viviano et al., 2017), is proposed in joint S-F researches to enrich the understandings beyond what is revealed by static FC alone. The large tract terminal on several cortical regions, which can be separable functionally, is mentioned as the reason for weak SC-FC correlation. To this end, using FC-derived seed regions instead of tract-derived is proposed (Fjell et al., 2017; Zimmermann et al., 2016) The correlation coefficient, which is used to obtain cortical networks, is discussed in (Romero-Garcia et al., 2014). Using absolute values of signed correlation instead of positive correlation values increases the level of S-F coupling. However, the interpretation of networks involving absolute correlations is complicated due to the use of anti-correlated values and the reduction of the small-worldness of such networks. Moreover, it is not highly suggested in age-related joint researches, in particular for the age range when transmodal-related improvements are more prevalent than unimodal changes. In addition, using graph metrics of SC, FC, and effective connectivity may provide valuable information in joint S-F studies.

Likewise, new techniques modeling characteristics of the brain more accurately may help to understand how the function and structure of the brain are associated with each other. For example, gneralized diffusion tensor imaging using higher order tensor statistics (GDTI-HOT) (Liu et al., 2010) and Neurite orientation dispersion and density imaging (NODDI) (Hui Zhang et al., 2012) provide a sophisticated model of diffusion MRI. These techniques are sensitive to microstructure changes and have been used as a decent interpretation in areas of complex axonal or dendritic architecture. Additionally, improving fiber-tracking methods to extract fine fibers and SC is also another important issue. There exist different challenges for fiber tracking (Jeurissen et al., 2019) like ambiguous local geometries, tracking fibers near the cortex, and angular resolution limits. The effect of fine fiber detection would be more apparent, considering the relation between function and gyri-gyri and sulci-gyri connections (Deng et al., 2014). In the last decade, improving the fiber tracking methods has received growing attention using the state-of-the-art of imaging techniques and computational approaches (Dell’Acqua & Tournier, 2019; Hongliang Zhang et al., 2013).

In conclusion, joint S-F biomarkers are supposed to improve brain temporal atlases and help to understand brain performance and its development. However, we are still far away from comprehensive and informative atlases that can show the maturation of structure and function of the brain, and help to predict disorders before behavioral and clinical symptoms appearance. Improving data, informative biomarkers, and techniques discussed in this section are crucial to progress in this area.

## Appendix A

**Table A1** through **Table A4** contain all abbreviations used throughout this paper and their meaning.

**Table A1.** List of abbreviations (A-F)

| A |  |
| --- | --- |
| ACC | Auditory fibers of the corpus callosum |
| AEC | Amplitude envelope correlation |
| AF | Arcuate fasciculus |
| AR | Acoustic radiations |
| C |  |
| CAEPs | Cortical auditory evoked potentials |
| D |  |
| DMN | Default mode network |
| DTI | Diffusion tensor imaging |
| DWI | Diffusion weighted imaging |
| E |  |
| EEG | Electroencephalography |
| ERN | Error-related negativity |
| F |  |
| FA | Fractional anisotropy |
| FC | Functional connectivity |
| FM | Functional modality |
| FNC | Functional network connectivity |
| fNIRS | Functional near-infrared spectroscopy |
| FPN | Frontoparietal network |

**Table A2.** List of abbreviations (G-L)

| G |  |
| --- | --- |
| GABA | Gamma aminobutyric acid |
| GAM | Generalized additive model |
| GDTI-HOT | Generalized diffusion tensor imaging using higher order tensor statistics |
| GFP | Global field power |
| GM | Gray matter |
| GMV | Gray matter volume |
| GPL | Global path length |
| GW | Gestational week |
| H |  |
| HCP | Human connectome project |
| HRA | Hierarchical regression analysis |
| I |  |
| IFG | Inferior frontal gyrus |
| ipsiL | Ipsilateral |
| L |  |
| LC | Local clustering |

**Table A3.** List of abbreviations (M-R)

| M |  |
| --- | --- |
| M | Month(s) |
| m | Male subject |
| MEG | Magnetoencephalography |
| MOG | Middle occipital gyrus |
| mPFC | Medial prefrontal cortex |
| MRI | Magnetic resonance imaging |
| mSTG | Middle superior temporal gyrus |
| MTL | Medial temporal lobe |
| N |  |
| NODDI | Neurite orientation dispersion and density imaging |
| P |  |
| PCC | Posterior cingulate cortex |
| Pe | Error positivity |
| PET | Positron emission tomography |
| PSD | Power spectral density |
| pSTG | Posterior superior temporal gyrus |
| R |  |
| rlPFC | Rostrolateral prefrontal cortex |
| ROI | Region of interest |
| RS | Resting-state |
| RSNs | Resting-state networks |

**Table A4.** List of abbreviations (S-Z)

| S |  |
| --- | --- |
| S-F | Structural and functional |
| SC | Structural connectivity |
| SCNs | Structural covariance networks |
| SIM | Structural imaging modality |
| SL | Synchronization likelihood |
| SN | Salience network |
| SNC | Structural network correlation |
| STG | Superior temporal gyrus |
| T |  |
| TMS | Transcranial magnetic stimulation |
| TPJ | Temporoparietal junction |
| V |  |
| VEPs | Visual evoked potentials |
| VEFs | Visual stimulus-evoked fields |
| W |  |
| WM | White matter |
| WMV | White matter volume |
| Y |  |
| Y | Year(s) |

## Declarations

### Funding

Not applicable

### Conflicts of interest/Competing interests

Not applicable

### Availability of data and material

Not applicable

### Code availability

Not applicable

### Authors’ contributions

Hamid Abrishami Moghaddam had the initial idea for this article, which was completed and organized by Roxana Namiranian and Sahar Rahimi. Roxana Namiranian and Sahar Rahimi performed the literature search. Sahar Rahimi and Roxana Namiranian drafted and all the authors critically revised the work.

### Ethics approval

Not applicable

### Consent to participate

Not applicable

### Consent for publication

Not applicable

## Acknowledgment

We thank Prof. Fabrice Wallois for his valuable suggestions on the initial idea of this paper. We would like also to thank Dr. Farveh Daneshvarfard for her comments on the initial phase of research, and Dr. Shirin Mavandadi for her comments on **Fig. 2**. We would particularly like to thank Cognitive Sciences and Technologies Council (COGC), Iran in the framework of Neurobiom project, for their support and guidance.

## Notes

### Competing Interest Statement

The authors have declared no competing interest.

